# In *Lyl1*^*-/-*^ mice, adipose stem cell vascular niche impairment leads to premature development of fat tissues

**DOI:** 10.1101/2020.04.24.059683

**Authors:** Abid Hussain, Leila El Kebriti, Virginie Deleuze, Yaël Glasson, Nelly Pirot, Danièle Mathieu, Valérie Pinet

## Abstract

Lymphoblastic leukemia-derived sequence 1 *(Lyl1)* encodes a hematopoietic- and endothelial-specific transcriptional factor. *Lyl1*-deficient mice are viable, but they display mild hematopoietic and vascular defects. Here, we report that young *Lyl1*^*-/-*^ mice exhibit transient obesity associated with general expansion of adipose tissues and unrelated to food intake. The increased fat tissue development in *Lyl1*^*-/-*^ mice resulted from an earlier adipocyte differentiation of adipose stem cells (ASCs) through non-cell autonomous mechanisms. Specifically, we found that in *Lyl1*^*-/-*^ mice, the vascular structures of adipose tissues are unstable, more prone to angiogenesis and, consequently, cannot maintain adipose progenitors in the niche vessel wall. Together, our data show that in *Lyl1*^*-/-*^ mice, the impaired vascular compartment of the adipose niche promotes uncontrolled ASC activation and differentiation, leading to early adipocyte expansion and premature depletion of ASCs. Our study highlights the major structural role of the adipose tissue vascular niche in coordinating stem cell self-renewal and differentiation into adipocytes.

## INTRODUCTION

Endothelial cells (ECs) are the major component of the vascular network that spreads into every organ of the body. Besides the conventional role of blood vessels in the transport of gases, nutrients, waste products and cells, ECs have been functionally linked to a wide range of physiological and pathological processes, including barrier formation, selective transport, scavenging, thrombosis, wound healing and inflammation. In the last decade, it has been shown that ECs are crucial regulators of tissue morphogenesis through secretion of growth factors and presentation of molecular signals that act on neighboring cell populations in an angiocrine fashion [1-3]. Moreover, blood vessels provide protective and supporting niche microenvironments for multiple adult stem and progenitor cells, such as neural [4, 5], muscle [6] and hematopoietic stem cells [7-10] as well as hepatic [11, 12] and adipocyte progenitors [13]. New vessel formation includes different sequential steps: basement membrane remodeling, perivascular cell detachment, EC proliferation and alignment, lumen formation, vessel maturation with adherens junction closure and finally pericyte coverage that stabilizes the vascular structures. Most of the mechanisms involved in the latest steps of vessel maturation contribute to the maintenance of mature vessel integrity [14].

The transcription factor lymphoblastic leukemia-derived sequence 1 (LYL1) is a member of the basic helix loop helix (bHLH) family. Its expression is restricted to hematopoietic [15] and endothelial [16] cell lineages both during embryonic life and adulthood. *Lyl1*-deficient (*Lyl1*^*-/-*^) mice develop normally without gross histological abnormalities. However, others and we highlighted the presence of hematopoietic and vascular system defects. Specifically, disruption of LYL1 activity in mice impairs the long-term hematopoietic reconstitution capacity and maintenance of early T lineage progenitors and partially blocks B lymphocyte differentiation [17, 18]. More recent studies from Chiu *at al.* demonstrated that *Lyl1* can maintain primitive erythropoiesis and compensate for loss of Scl (a LYL1 co-member of bHLH family) in megakaryopoiesis [19, 20]. LYL1 is also required for the maturation and stabilization of endothelial adherens junctions of newly formed vessels [16, 21]. Consequently, blood vessels in the lung of young adult *Lyl1*^*-/-*^ mice cannot form a functional endothelial barrier, leading to vascular leakiness [21]. In *Lyl1*^*-/-*^ mice grafted with cancer cells, tumor vessels are highly angiogenic, fully immature and poorly covered by mural cells [16].

During these previous studies, we noticed that young adult *Lyl1*^-/-^ mice were often overweight with generalized increase of fat mass. While obesity classically results from white adipose tissue (WAT) expansion, *Lyl1* absence affected all adipose depots: brown adipose tissue (BAT) as well as subcutaneous and visceral WAT. Here, we show that the accelerated development of the three adipose tissues in *Lyl1*^*-/-*^ mice results from non-cell autonomous mechanisms. Specifically, we identified premature development of vascular structures in *Lyl1*^*-/-*^ juvenile adipose organs triggering faster mature adipocyte formation. Together, our data show that the immature and unstable vascular structures of *Lyl1*^*-/-*^ adipose tissues prevent adipocyte stem cell maintenance in the niche vessel walls, thereby accelerating their differentiation into adipocytes and causing premature stem cell depletion.

## MATERIALS and METHODS

### Mice

*Lyl1*^*-/-*^ mice have been previously described [17]. Mice were housed in temperature-controlled ventilated cages (20-22°C) with a 12h light-dark cycle and maintained in pathogen-free conditions in the institute animal facility. All experiments were conducted by authorized personnel, in accordance with the European Union directive n°2010/63/EU and approved by the Languedoc-Roussillon Animal Care and Use Committee (agreement number CEEA-LR 12061). Mice genotypes were determined by PCR assay of tail DNA, as described elsewhere [16]. Male mice were preferentially used to avoid interference by the female hormonal cycle. Mice were anesthetized by isofluorane inhalation, sacrificed and BAT and WAT collected, weighted and used for gene expression analyses or fixed overnight in neutral buffered formalin (4% formaldehyde), dehydrated and embedded in paraffin for histological analyses. Body composition (fat and lean mass) was assessed by nuclear magnetic resonance using an EchoMRI 3-in-1™ analyzer (Service de phénotypage de la plateforme Anexplo, Toulouse, France).

### Adipocyte size and beige fat area

Four µm-thick tissue sections of paraffin-embedded WT and *Lyl1*^*-/-*^ adipose tissue were stained with hematoxylin-eosin and visualized with a NanoZoomer slide scanner controlled by the NDP.view software. For BAT and eWAT, lipid droplet size in at least 5000 adipocytes was measured on 3-5 different sections using the ImageJ software with the special MRI-plugging “Adipocyte_tools” (designed by V. Baecker from the Montpellier Resources Imaging facility). In eWAT tissue sections, objects that fell below the area of 350µm^2^ were removed because they could be a mixture of stromal vascular cells [22]. Beige fat zone areas in ingWAT were measured with the NDP.view software and indicated as percentage relative to the total surface of the ingWAT section.

### SVF isolation and culture

SVF cells were isolated from BAT and ingWAT of 8-day-old puppies and 12-week-old males, respectively. Fat pads were excised, finely cut with scissors and incubated in digestion medium (PBS with 2% BSA, 1M CaCl_2_, 94U/ml dispase II and 100mg/ml collagenase D, Roche Life Science, France) at 37°C for 30-40min. Floating adipocytes were separated from the SVF by centrifugation at 500g for 5min. SVF cells were sequentially filtered through 100- and 40-µm filters and then plated in Petri dishes with DMEM-F12 and Glutamax (Thermo Fischer Scientific, France) supplemented with 10% FBS and 5ng/ml bFGF. Culture medium was changed every two days.

### Flow cytometry and cell sorting

SVF cells prepared as described above were stained with anti-CD45 PerCP (BD Biosciences; 557235), -CD31 PE-Cy7 (BD Biosciences; 561410), -CD34 eFlour 660 (eBioscience, 50-0341), -SCA-1 APC-Cy7 (BD Biosciences, 560654), -CD24 FITC (eBioscience, 11-0241) and -CD140a PE (eBioscience, 12-1401) antibodies at 4°C for 30min. Samples were then washed and centrifuged at 300g for 5min. ASCs and pre-adipocytes were analyzed with a BD FACS-Quanto II or sorted with a BD FACS Aria and data analyses were performed using the BD FACS Diva software.

### SVP isolation, culture and staining

eWAT and ingWAT from 12-week-old male mice were cut in 5-10 pieces and incubated with 1 and 2mg/ml, respectively, of collagenase at 37°C for 2h. Mixtures were passed through a 300µm mesh to remove big fragments and then through a 30µm mesh to remove single cells. The remaining SVP on the 30µm mesh were washed off in DMEM-F12 with Glutamax (Thermo Fischer Scientific, France) supplemented with 10% FBS and seeded on coverslips coated with 2% gelatin. Microtubules were allowed to grow from SVP in DMEM-F12+10% FBS for 5 days before staining with an anti-CD31 antibody (BD Biosciences, 557355). Images of microtubule outgrow from SVP were acquired with a Leica SP5-SMD confocal microscope. The angiogenic response was determined in each individual SVP sample by measuring the length of the growing microtubules with the Image J software. Results are presented as the mean ± standard error of the mean (SEM) of the tubule length and analyzed with the Mann-Whitney test and GraphPad Prism 5.0 (MacKiev).

### Statistical analysis

All statistical analyses were performed with the GraphPad Prism 5.0 program (Mackiev software). After testing for data normality, the unpaired *t* test was used when variances were not significantly different and the unpaired *t* test with Welch’s correction when variances were significantly different. When Gaussian distribution was not assumed Mann-Whitney test was used.

See *Supplemental Materials* for Metabolic parameter survey, RNA preparation, mRNA expression analysis, Immunohistochemistry, adipogenic differentiation, clonogenicity test, Oil Red O staining, blood vessel immunofluorescence staining, extravasation of albumin-Evans blue within tissues and whole-mount confocal microscopy.

## RESULTS

### Young adult *Lyl1*^-/-^ mice are overweight and display an overall increase of fat mass due to early expansion of all adipose tissue

During our phenotypic analysis of *Lyl1*^-/-^ mice, we noticed that young adults were consistently overweight. Specifically, when compared with wild type (WT) littermates, *Lyl1*^-/-^ males showed a significantly higher body weight increase from week 8 to week 18 post-partum. This difference reached 14% at week 14 post-partum (Figure 1A). At week 22 post-partum, body weight reached a comparable plateau in both WT and *Lyl1*^*-/-*^ mice. Echo magnetic resonance imaging analysis of 10-week-old animals showed a significantly higher percentage of total fat mass in *Lyl1*^*-/-*^ than WT males, whereas the lean body mass fraction was comparable between groups (Figure 1B). Food intake monitoring for six weeks showed no significant difference between genotypes (Figure S1A).

**Figure 1.**
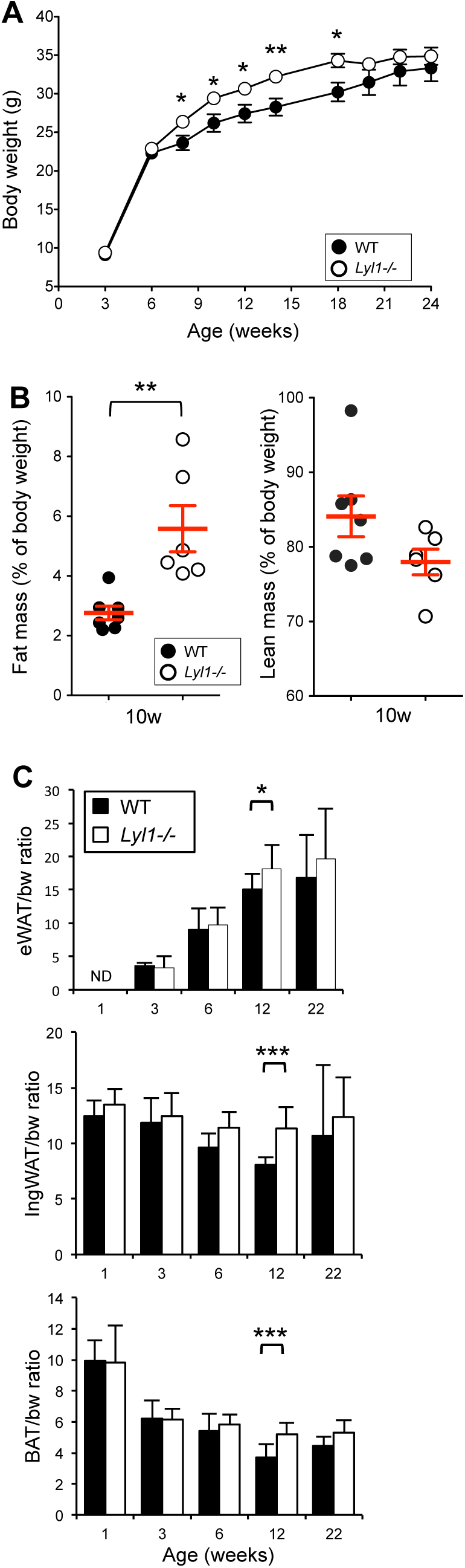
Young adult *Lyl1*^-/-^ mice, display an overall expansion of adipose tissue. **(A)** Body weight changes in 3 to 24-week-old wild type (WT) (n=6-12) and *Lyl1*^*-/-*^ (n=4-6) mice. **(B)** Body composition at 10 weeks of age was assessed by nuclear magnetic resonance using an EchoMRI whole body composition analyzer. Body fat and lean mass are indicated as percentage of body weight. **(C)** Epididymal WAT (eWAT), inguinal WAT (ingWAT) and BAT to body weight (bw) ratios were determined in 1-, 3-, 6-, 12- and 22-week-old WT (n=5-10) and *Lyl1*^*-/-*^ (n=5-10) mice. ND: Not Detected. Results are presented as the mean ± SD of adipose tissue mass (mg)/bw (g) (‰). \**P*<0.05; \*\**P*<0.01; \*\*\**P*<0.001.

There is two major adipose tissues: BAT that plays a central role in energy expenditure to produce heat, and WAT that is specialized in energy storage as fatty acids. As total fat mass increase is associated particularly with WAT expansion, we examined epididymal WAT (eWAT) and inguinal WAT (ingWAT), as examples of visceral and subcutaneous WAT respectively, at different ages. Compared with WT mice, the ratios of eWAT and ingWAT to body weight were significantly higher in 12-week-old *Lyl1*^*-/-*^ mice (Figure. 1C). BAT also was significantly increased in *Lyl1*^*-/-*^ mice at 12 weeks of age (Figure. 1C).

We assessed whether overweight was associated with metabolic syndrome. *Lyl1*^*-/-*^ mice did not show any change in glucose tolerance, insulin secretion in response to glucose load, insulin tolerance and hepatic glucose production (assessed with the pyruvate tolerance test) compared with WT animals (Figure S1B). To investigate whether overweight in *Lyl1*^*-/-*^ mice was caused by deregulated energy homeostasis, we housed adult males in individual metabolic cages. *Lyl1*^*-/-*^ and WT mice showed similar energy expenditure and globally comparable exchanges of oxygen and carbon dioxide (RER, respiratory exchange ratio). However, more precisely, the non-reduction of RER at the end of the light cycle for the *Lyl1*^*-/-*^ mice (when animals were fasting) suggests a defect in lipid oxidation usually supported by BAT (Figure S1C).

### Increased adiposity of all fat tissues of young Lyl1-/- mice

To determine whether fat pad mass increase was caused by enhanced adipocyte growth (hypertrophy) and/or number (hyperplasia), we analyzed paraffin-embedded adipose tissue sections stained with hematoxylin-eosin. Lipid droplets were bigger in *Lyl1*^*-/-*^ than in WT eWAT samples at 12 weeks, but not at 3, 6 and 22 weeks of age (Figure 2A). Similarly, histological analysis of BAT sections of 3-, 6-, 12- and 22-week-old mice housed at 21-22°C and fed the classical diet showed that lipid droplets were larger in 12-week-old *Lyl1*^*-/-*^ than WT BAT (Figure 2B), classically called a whitening of BAT. In 22-week-old mice, lipid droplets were relatively heterogeneous in size and did not significantly differ between genotypes. Likewise, analysis of the number of brown-like adipocytes by measuring the beige zone areas in ingWAT sections showed marked reduction of beige fat zones associated with a global enlargement of lipid droplets in 12-week-old *Lyl1*^*-/-*^ ingWAT compared with WT samples (Figure 2C). In agreement, the expression of beige adipocyte-specific genes, such as *Tmem26* and *Cd137*, was reduced in 12-week-old *Lyl1*^*-/-*^ ingWAT compared with WT tissue as usually observed in older animals (data not shown). Interestingly, beige zone areas were increased in 3-week-old *Lyl1*^*-/-*^ ingWAT compared with WT, suggesting an early functionality of this tissue in *Lyl1*^*-/-*^ mice. The contribution of hyperplasia to fat mass expansion could be excluded since immuno-histochemical analysis using the proliferative marker Ki67 on tissue sections of eWAT (Figure S2), BAT and ingWAT (data not shown) of 6 and 12-week-old animals did not show any difference between WT and *Lyl1*^*-/-*^ mice. Together, these data indicate that weight increase in *Lyl1*^*-/-*^ mice is due to fat mass expansion and global enlargement of lipid droplets in all adipose tissues.

**Figure 2.**
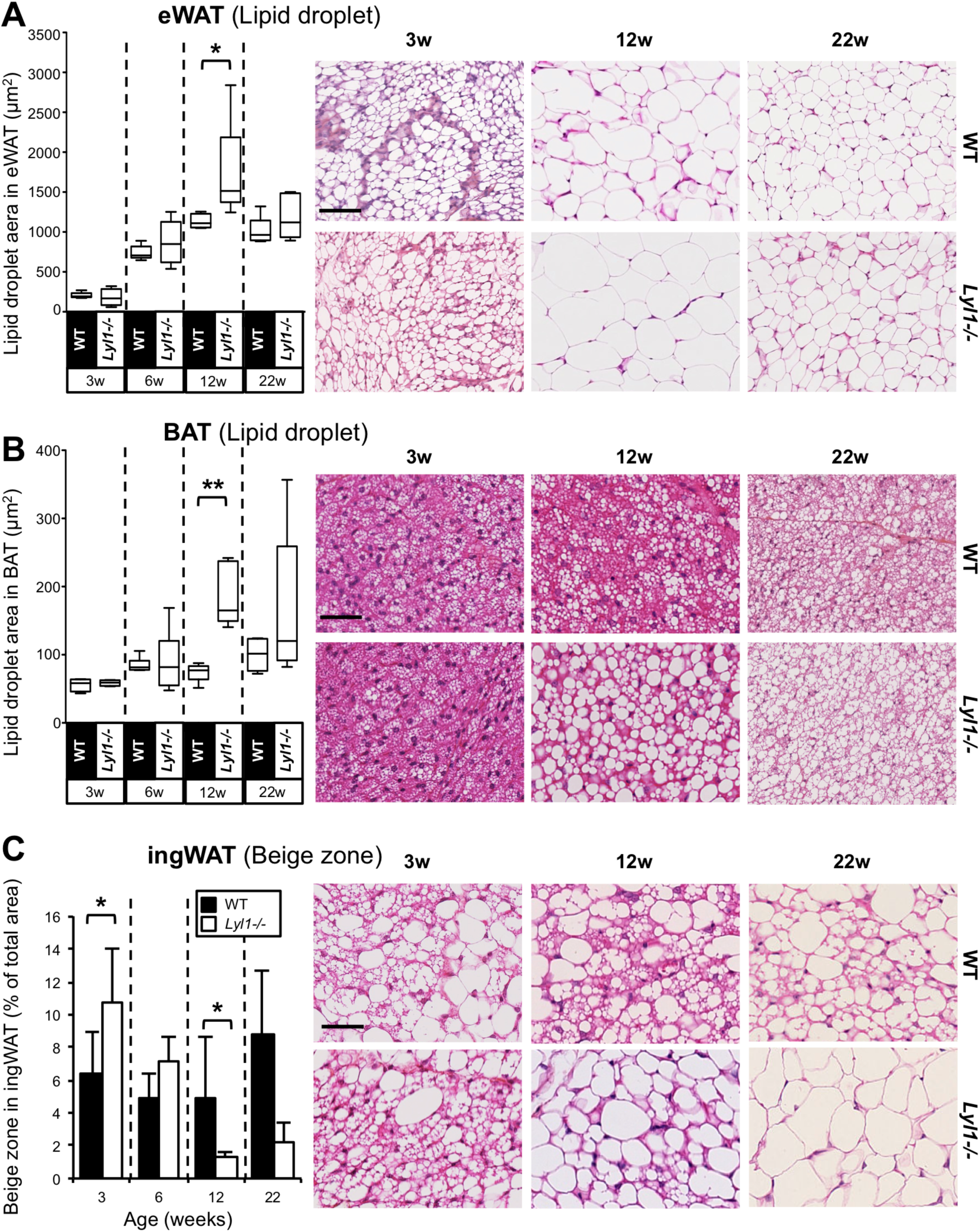
Increased adiposity in all adipose tissues of young adult Lyl1^-/-^ mice. **Left panels**: Lipid droplet size in eWAT (A) and BAT (B) was measured as described in the Materials and Methods section. Results are presented as the median lipid droplet area (whiskers = min to max) and analyzed with the Mann-Whitney test. \**P*<0.05, \*\**P*<0.01. The beige fat zone area in IngWAT (C) was measured with the NDP.view software and relative to the total surface of the ingWAT section. Results are presented as the mean ± SD of beige fat zone percentage and analyzed with the Mann-Whitney test. **Right panels**: Paraffin-embedded adipose tissue sections from WT and *Lyl1*^*-/-*^ mice (n=5-8 per age and genotype) were stained with hematoxylin-eosin and images visualized with a NanoZoomer slide scanner controlled by the NDP.view software. Scale bar: 50µm.

### Early activation of the thermogenic potential in BAT and premature aging features in eWAT of young adult *Lyl1*^*-/-*^ mice

Due to the concomitant BAT whitening and decrease in beige fat zones in ingWAT from 12-week-old *Lyl1*^*-/-*^ mice (two signs of aging), we asked whether LYL1 absence alters the thermogenic potential. Immunostaining of BAT sections showed a significant decrease of UCP1, a protein that mediates heat generation, in BAT of 12-week-old *Lyl1*^*-/-*^ (Figure 3A) compared with WT mice, in agreement with BAT whitening. Moreover, *Cidea* mRNA expression was higher in BAT samples from 12-week-old *Lyl1*^*-/-*^ than WT mice (Figure 3B). This is consistent with lipid droplet enlargement (see Figure 2B), given CIDE-A role in the control of lipid storage and droplet enlargement [23]. Conversely, in very young animals (6 weeks of age and younger), UCP1 expression was significantly higher in *Lyl1*^*-/-*^ BAT than in controls, suggesting that the thermogenic program might be active earlier in *Lyl1*^-/-^ mice. This was confirmed by the increase of beige zone area revealed as soon as 3 weeks in *Lyl1*^*-/-*^ ingWAT (see Figure 2C). Similarly, *Adrβ3* expression was higher in 1- and 3-week-old *Lyl1*^*-/-*^ than WT BAT, suggesting a stronger potential response to sympathetic β3-adrenergic stimulation (Figure 3B). The higher *Cidea* expression in *Lyl1*^*-/-*^ than WT puppies (1-week-old; Figure 3B) suggested early fat browning, a process involved in the acquisition of the multilocular morphology characteristic of functional brown adipocytes [24]. To evaluate the consequence of BAT and ingWAT whitening on tissue function, we implanted temperature-sensitive transmitters intraperitoneally in 12-week-old mice that were housed in individual metabolic cages at 22-24°C for 3 days. Continuous monitoring of the internal temperature for 2 days showed significantly lower temperature in *Lyl1*^-/-^ mice than in WT controls (Figure 3C). Given the major role of BAT and subcutaneous WAT in thermogenesis, we assessed the behavior of 12-week-old WT and *Lyl1*^*-/-*^ mice upon exposure to 9°C in individual metabolic cages for 24hrs (Figure S3). Even if the internal body temperature was lower in *Lyl1*^-/-^ mice than in WT controls this was not significant, indicating that Lyl1^-/-^ mice can respond to cold stress almost as efficiently as WT mice.

**Figure 3.**
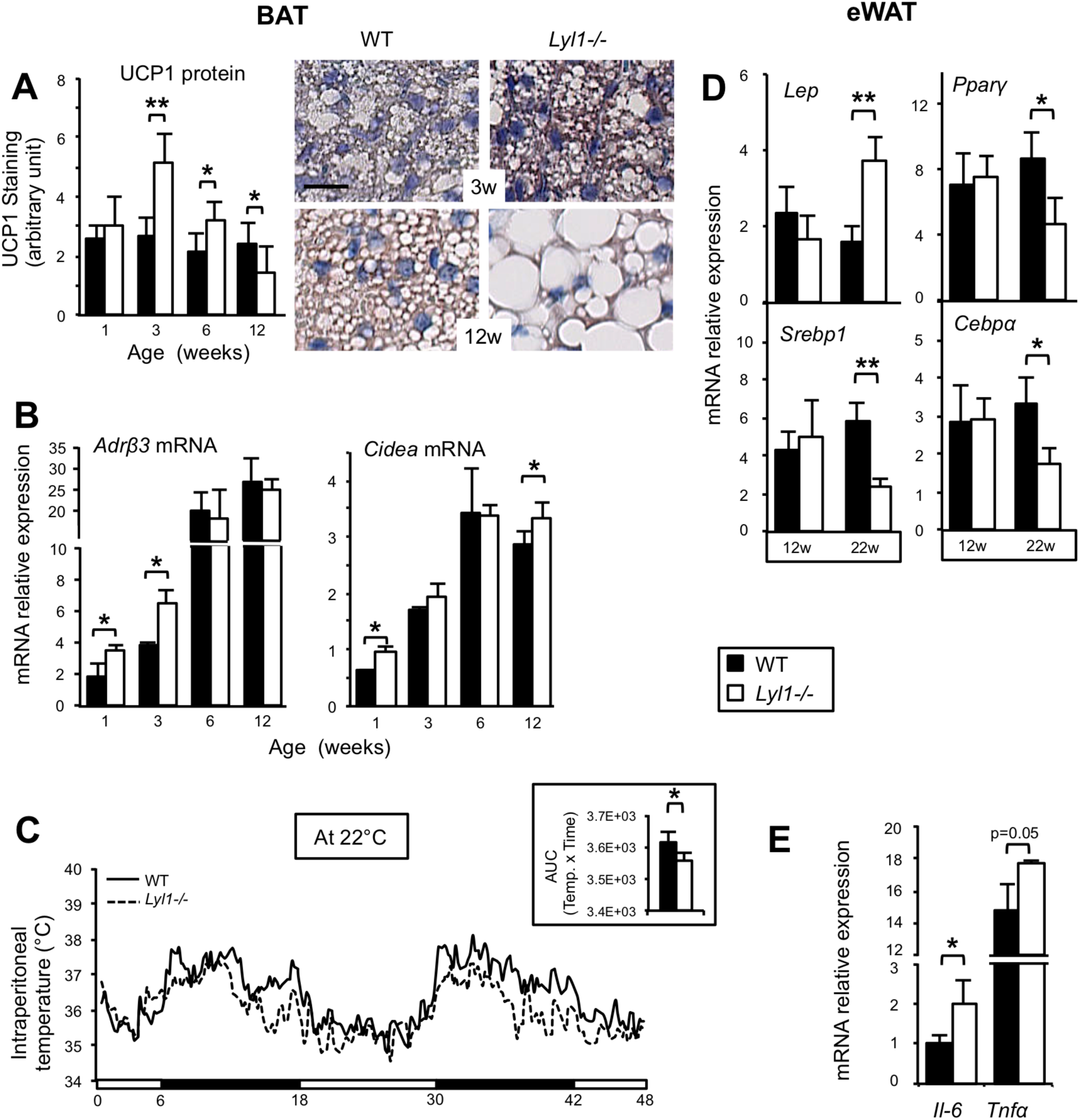
Early activation of the thermogenic potential in BAT and premature aging features in eWAT of young adult Lyl1^-/-^ mice. **(A)** Quantification with Aperio ImageScope of UCP1 protein immunostaining in paraffin-embedded BAT sections from WT (n=3-7 per age) and *Lyl1*^*-/-*^ (n=4-8 per age) mice. Results are relative to the total surface of the BAT section. BAT tissue sections from WT and *Lyl1*^*-/-*^ mice were immunostained for UCP1 protein and images visualized with a NanoZoomer slide scanner controlled by the NDP.view software. Scale bar: 20µm. **(B)** Total RNA was extracted from BAT samples from 1-, 3-, 6- and 12-week-old WT and *Lyl1*^*-/-*^ mice (n=3-6 per group). Expression of *Adrβ3* and *Cidea* was quantified by qPCR and normalized to *Actβ*. **(C)** Intraperitoneal temperature monitoring in 12-week-old mice housed individually in metabolic cages with a temperature of 22°C for two days (3 mice for each genotype). *Inset*: The area under the curve (AUC) was calculated using the trapezoidal rule. **(D)** Total RNA was extracted from eWAT samples from 12- and 22-week-old WT and *Lyl1*^*-/-*^ mice (n=6-7 per group). Expression of *Lep, Srebf1, Pparγ* and *C/ebpα* was quantified by qPCR and normalized to *Actβ*. **(E)** Expression of the genes encoding the pro-inflammatory cytokines IL-6 and TNFα was quantified by qPCR in total RNA from 22-week-old WT and *Lyl1*^*-/-*^ eWAT samples (n=3 per group) and normalized to *Actβ* expression. \**P*<0.05; \*\**P*<0.01.

In 22-week-old mice, lipid droplet size in eWAT samples was comparable between genotypes (see Figure 2A), suggesting that accumulation of large adipocytes in eWAT declines in older *Lyl1*^*-/-*^ male*s*. Such a reduction of adipocyte size is typically observed in white adipose tissues of aged mice [25, 26]. High levels of circulating LEPTIN in plasma and high *leptin* gene expression in adipocyte tissues are also correlated with aging in both rodents and humans [27-30]. Therefore, we analyzed the mRNA expression of *leptin* (*Lep*) and *Srebp1*, its downstream target encoding the transcription factor SREBP1 that controls lipogenic genes in eWAT. Compared with WT animals, *Lep* mRNA expression was increased by more than 2-fold in 22-week-old *Lyl1*^*-/-*^ mice and *Srebp1* was downregulated (Figure 3D). As a consequence, the mRNA levels of the lipogenic genes *Acaca, Fasn* and *Scd1* were also reduced in 22-week-old *Lyl1*^*-/-*^ eWAT (data not shown). The downregulation of *C/ebpα* and *Pparγ*, two adipogenic genes, in 22-week-old *Lyl1*^*-/-*^ eWAT (Figure 3D) further supports the hypothesis of the premature eWAT maturation in *Lyl1*^*-/-*^ mice. As upregulation of pro-inflammatory cytokines is classically observed in adipose tissues of aged mice [31], we also evaluated *Il-6* and *Tnfα* expression in 22-week-old eWAT. Compared with WT samples, *Il-6* was significantly upregulated in *Lyl1*^*-/-*^ eWAT, but not *Tnfα*, although its expression tended to be higher in *Lyl1*^*-/-*^ eWAT (Figure 3E).

These data indicate that both WAT and BAT develop and mature earlier in *Lyl1*^*-/-*^ mice, leading to early activation of the thermogenic program of BAT and beige adipocytes and acceleration of aging-like processes, *i.e.* BAT whitening and brown-like adipocytes within ingWAT classically observed in older adult mice [30, 32].

### Early development of adipose tissues leads to premature decline of the adipogenic potential of the stromal vascular fraction of young adult *Lyl1*^*-/-*^ mice

The differentiation of immature progenitor cells into fully mature adipocytes involves the sequential activation of transcriptional programs [33]. To investigate the mechanisms leading to the adipocyte faster maturation in young *Lyl1*^*-/-*^ mice, we assessed the expression of genes encoding transcription factors involved in adipogenic processes in BAT and WAT at different ages (Figure 4A). Compared with WT, the expression of the early-acting genes *Zfp423* and *Klf4* decreased earlier in *Lyl1*^*-/-*^ BAT and ingWAT. Accordingly, the late-acting genes *Pparγ, C/ebpα* and *Srebp1* were activated earlier in *Lyl1*^*-/-*^ than WT BAT and ingWAT. In addition, the expression of *C/ebpα* and *Srebp1*, normally down-regulated upon aging, was prematurely reduced (12 weeks post-partum) in *Lyl1*^*-/-*^ BAT and ingWAT. Together, these data show that the adipogenic differentiation program is accelerated in *Lyl1*^*-/-*^ mice.

**Figure 4.**
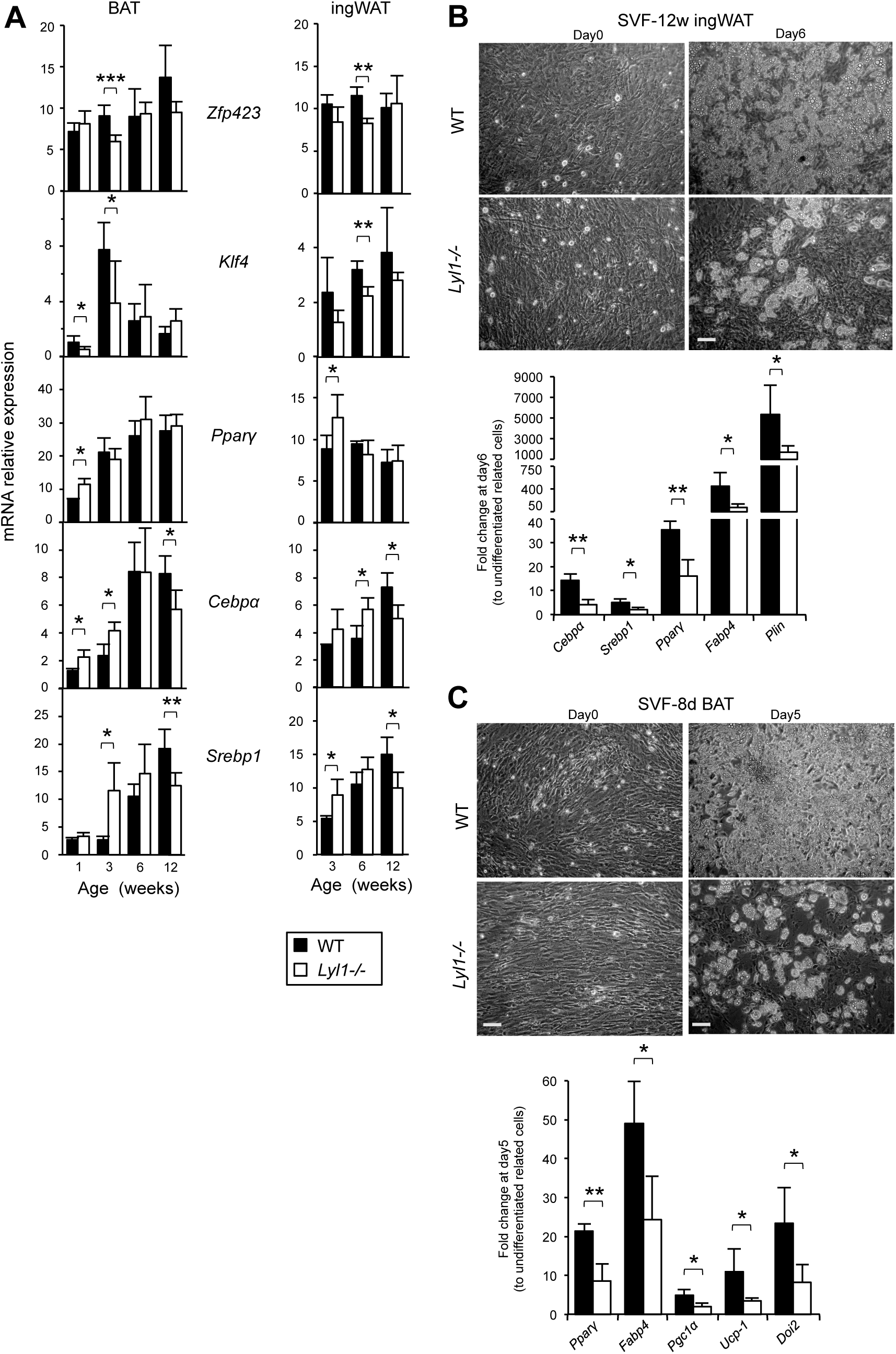
Premature onset of adipogenic differentiation in *Lyl1*^*-/-*^ BAT and ingWAT results in reduced adipogenic potential of stromal vascular fractions (SVFs). **(A)** Total RNA was extracted from BAT and ingWAT samples collected from 1-, 3-, 6- and 12-week-old WT and *Lyl1*^*-/-*^ mice (n=4-7). Expression of adipocyte differentiation genes (*Zfp423, Klf4, C/ebpα, Pparγ* and *Srebf1*), normalized to *Actβ*, was quantified by qPCR. **(B-C)** SVFs isolated from ingWAT samples (B) of 12-week-old mice (n=3 per group) or BAT (C) of 8-day-old mice (n=3 per group) were differentiated into adipocytes for 6 or 5 days, respectively, and then expression of adipogenic transcription factors (*C/ebpα, Pparγ* and *Srebf1*) and adipocyte-related markers (*Fabp4, Pgc1α, Ucp1, Dio2* and *Plin*) was quantified by qPCR and normalized to *36B4*. Upper panels: Representative images of SVFs from ingWAT at day0 and day6 of adipocyte differentiation (B) and of SVFs from BAT at day0 and day5 of differentiation (C). Scale bar: 100µm. \**P*<0.05; \*\**P*<0.01; *** P<0.001.

Adipocyte progenitors reside in the stromal vascular fraction (SVF) together with ECs, pericytes, fibroblasts and immune cells. To compare the adipogenic potential of fat tissues of young adult WT and *Lyl1*^*-/-*^ males, we cultured equal numbers of SVF cells from 12-week-old ingWAT samples to confluence before induction of adipocyte differentiation. After 6 days in adipogenic differentiation medium, we observed fewer mature adipocytes in *Lyl1*^*-/-*^ than in WT ingWAT-SVF cultures (Figure 4B). Expression analysis of genes encoding pro-adipogenic transcription factors (*C/ebpα, Srebp1* and *Pparγ*) and of mature adipocyte markers (*Fabp4* and *Plin*) confirmed the lower adipogenic potential of *Lyl1*^*-/-*^ SVF. Similarly, upon adipogenic differentiation, we observed fewer mature adipocytes in SVF cultures from BAT of 8-day-old *Lyl1*^*-/-*^ than WT puppies, in agreement with the lower expression of *Pparγ* and *Fabp4* as well as of BAT-specific markers (*Pgc1α, Dio2* and *Ucp1*) (Figure 4C). We then compared the capacity of progenitors contained in *WT* and *Lyl1*^*-/-*^ ingWAT-SVF to generate clones that can differentiate into mature adipocytes (Figure S4) by clonogenic assay. *Lyl1*^*-/-*^ ingWAT-SVF cultures exhibited less hematoxylin-positive clones than WT controls (64.5±17.7 versus 125.5±3.5) and, among them, fewer could differentiate into Oil Red O-positive mature adipocytes (17% versus 45%).

These data indicate that SVF cells derived from *Lyl1*^*-/-*^ adipose tissues contains fewer immature progenitors that can produce mature adipocytes, leading to a global reduction of the adipogenic potential of ingWAT and BAT in young *Lyl1*^*-/-*^ mice.

### Adipocyte progenitors are less numerous in *Lyl1-/-* adipose tissue-SVF and poorly associated with vasculature

These findings prompted us to investigate how *Lyl1* might influence SVF adipogenic capacity and specifically the number of available functional immature progenitors. In subcutaneous WAT, two different cell progenitors (uncommitted adipocyte stem cell (ASC) and committed pre-adipocytes) have been identified based on the expression of specific markers [34]. Accordingly, by incubating SVF cells from ingWAT, but also BAT, samples of 3-, 6- and 12-week-old mice with antibodies against the surface markers CD31 (ECs), CD45 (hematopoietic cells), CD34 and SCA-1 (immature cells), CD140a and CD24 [34], we could isolate and quantify ASCs (CD45-CD31-CD34+ SCA-1+ CD140a+ CD24+) and pre-adipocytes (CD45-CD31-CD34+ SCA-1+ CD140a+ CD24-) (Figure 5A). The fraction of ASCs was significantly lower in ingWAT- and BAT-SVF from 6- and 12-week-old *Lyl1*^*-/-*^ than WT mice (Figure 5B, upper panel). Pre-adipocytes also were markedly reduced in ingWAT-SFV from 6-week-old *Lyl1*^-/-^ compared with WT mice (Figure 5B, lower panel). ASC visualization in WAT samples from 12-week-old mice by immunofluorescence showed the presence of several CD31-CD140a+ CD24+ ASCs (asterisks in Figure 5C) that were associated with CD31+ structures or in the tissue parenchyma in WT samples. Conversely, ASCs were rare in *Lyl1*^*-/-*^ eWAT samples.

**Figure 5.**
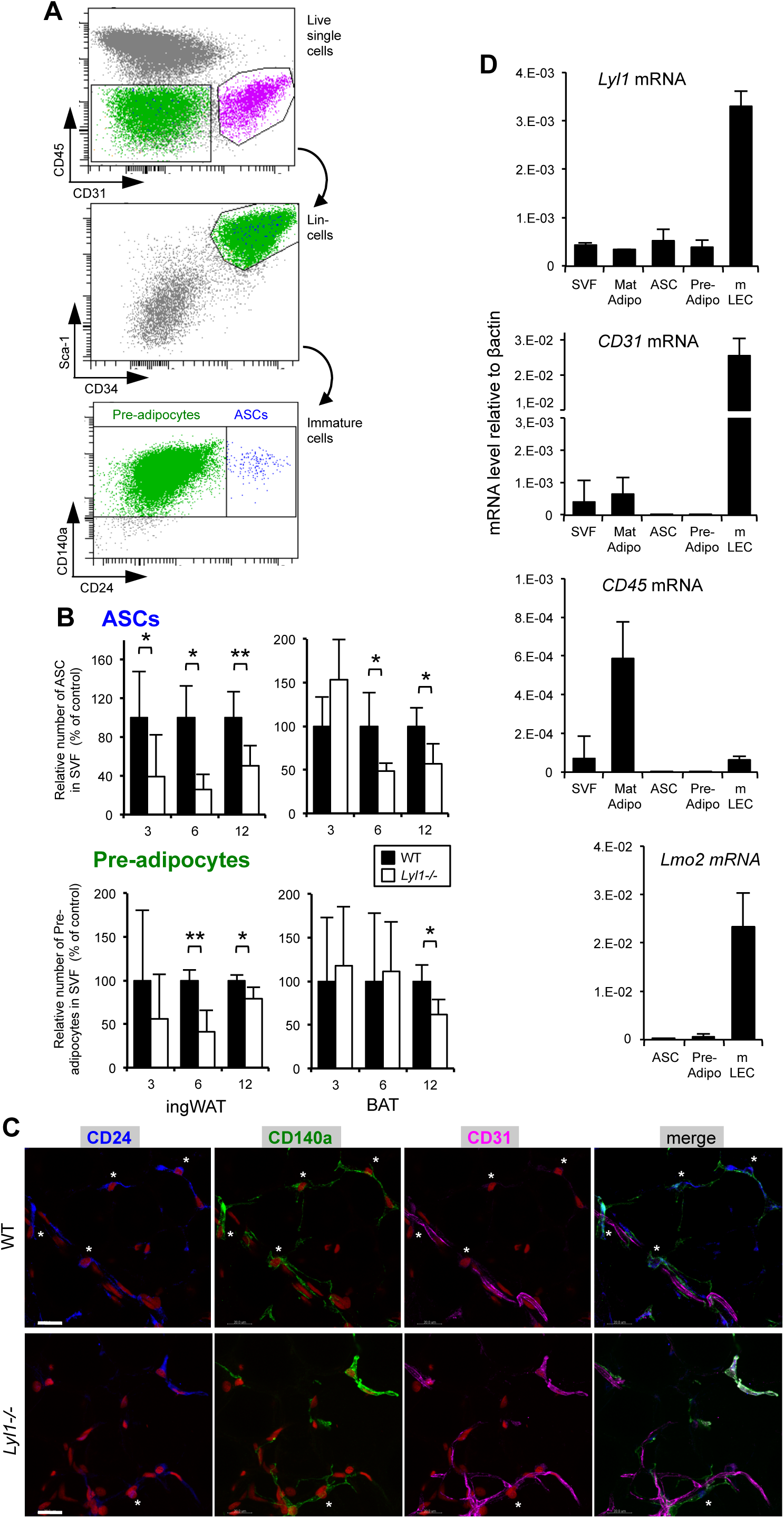
Adipose stem cells (ASCs) are less numerous ingWAT and BAT from in *Lyl1*^*-/-*^ mice and *Lyl1* is not expressed in the adipocyte lineage. **(A)** Dot plots showing FACS analysis of SVFs isolated from ingWAT of a 6-week-old WT mouse and the successive gating to isolate pre-adipocytes (CD140a+ CD24-, green) and ASCs (CD140a+ CD24+, blue). **(B)** ASCs and pre-adipocytes were quantified by flow cytometry in SVFs isolated from ingWAT and BAT of 3-, 6- and 12-week old WT and *Lyl1-/-* mice. Results are presented as the relative number of ASCs or pre-adipocytes in SVFs (i.e., number of cells divided by the mean number in the WT condition x 100). The fraction of ASCs and pre-adipocytes in adipose tissues varied from 0.05 to 0.20% and from 10 to 35% of live cells, respectively. n=3-6 mice/age/genotype. **(C)** ASCs were visualized in whole-mount 12-week-old eWAT as CD24+ CD140a+ CD31-cells (stars). Scale bar: 20µm. **(D)** Analysis of *Lyl1, Lmo2, CD31, CD45* and *Pparγ* mRNA expression in the indicated cell fractions and primary cells by RT-PCR. Total RNA was extracted from cells derived from 12-week-old WT mice (n=3). SVF was prepared from ingWAT, and mature adipocytes (Mat Adipo) were isolated from eWAT. ASCs and pre-adipocytes (Pre-Adipo) were FACS-sorted from ingWAT-SVF using specific markers, and mouse lung endothelial cells (mLECs) were prepared from dissociated lungs as previously described [21]. cDNAs were amplified in triplicate with specific mouse primers (Table S1) and normalized to Actβ; the means ± SD are shown.

### *Lyl1* is not expressed in the adipocyte lineage

We next wanted to determine the mechanisms by which *Lyl1* regulate the number of ASC and progenitor cells. We checked *Lyl1* expression in FACS-isolated ASCs and pre-adipocytes in comparison to mature adipocytes and endothelial cells (Figure 5D). *Lyl1* was expressed in SVF, in agreement with the presence of many CD31+ ECs and CD45+ hematopoietic cells, known to express *Lyl1. Lyl1* expression was very low in purified ASCs and pre-adipocytes (15.6% and 11.7% respectively), compared with purified mouse lung endothelial cells (mLECs). Moreover, ASCs and pre-adipocytes are unlikely to exhibit an active and functional LYL1 protein because they do not expressed *Lmo2*, a gene that encodes an obligate LYL1 partner in ECs and hematopoietic cells [35]. The weak *Lyl1* expression in the mature adipocyte fraction could be due to the faint contamination by *Lyl1*-expressing cells, such as ECs and hematopoietic cells. Furthermore, *Lyl1* was not expressed in committed 3T3-L1 pre-adipocytes at any stage of adipocyte differentiation (Figure S5A). Importantly, equal numbers of ASCs and pre-adipocytes isolated from ingWAT-SVF of 12-week-old WT and *Lyl1*^*-/-*^ mice reached confluence at the same time, suggesting comparable proliferative rates (data not shown). Moreover, when stimulated with the adipogenic cocktail, they underwent similar robust mature adipocyte differentiation (Figure S5B). This indicates that *Lyl1* absence has no effect on the intrinsic adipogenic potential of immature progenitors. Together, these data demonstrate that *Lyl1* exerts a non-cell autonomous function on adipocyte development.

Considering that Lyl1 is expressed in myeloid lineage and that obesity has been associated with accumulation of newly recruited macrophages in adipose tissue [36, 37], we assessed the presence of F4/80+ macrophages in eWAT and ingWAT of 6- and 12-week-old *Lyl1-/-* and WT mice. Compared to WT, *Lyl1-/-* adipose tissues did not show any increased accumulation of macrophages, either before the onset of overweight (6-week-old mice) or during the expansion of fat (12-week-old mice) (Figure S6).

### Blood vessels in *Lyl1*^*-/-*^ adipose tissues are immature and prone to angiogenesis

Lineage tracing studies have shown that within fat tissues, the capillary vascular wall is the niche of adipocyte progenitors [38, 39]. As we previously reported that *Lyl1* is required for the maturation and stabilization of newly formed vessels [16, 21], we now investigated whether *Lyl1* absence also affects the vascular compartment of the adipose tissue niche.

To evaluate vessel coverage in BAT vascular structures, we double stained BAT cryosections with anti-CD31 (ECs) and anti-NG2 (pericytes) antibodies. Quantification of the NG2-positive area relative to the CD31-positive surface showed that blood vessels in BAT were 66% less covered by pericytes in *Lyl1*^*-/-*^ than WT samples (Figure 6A), leading to a disorganized and unstable architecture of blood vessels (Figure S7A). This poor coverage of blood vessel by pericytes was confirmed by the use of a second marker for pericytes, PDGFRβ (Figure S7B). As VE-CADHERIN is a major actor in establishing functional endothelial barriers and in maintaining their integrity [40], we analyzed its localization in ingWAT endothelium of 12-week-old mice previously injected *in vivo* with FITC-labeled lectin to visualize blood vessels. VE-CADHERIN staining at cell-cell junctions was significantly decreased in *Lyl1*^*-/-*^ (by about 50%) compared with WT ingWAT samples (Figure 6B). The poor vessel coverage by mural cells and the defective recruitment of VE-CADHERIN at EC junctions strongly suggested that the vascular barrier integrity was affected in *Lyl1*^*-/-*^ adipose tissue, as observed in lungs of young *Lyl1*^*-/-*^ mice [21]. Evans blue dye extravasation was increased in both ingWAT and BAT in 12-week-old *Lyl1*^*-/-*^ mice compared with WT animals (2.3 and 1.4-fold higher in ingWAT and BAT, respectively), confirming the vessel structure leakiness and instability in *Lyl1*^*-/-*^ mice (Figure 6C).

**Figure 6.**
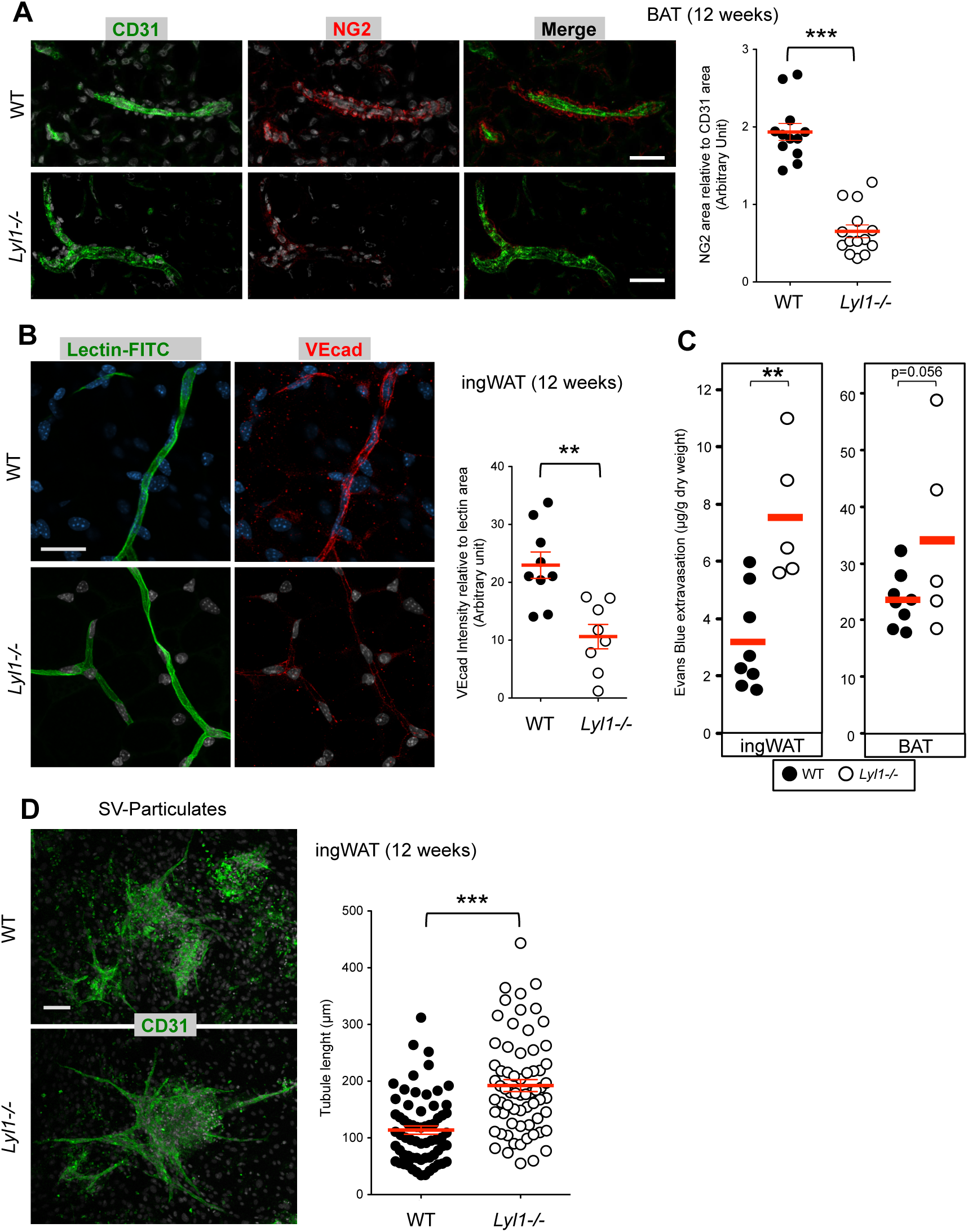
The vascular niche of adipocyte precursors is altered in *Lyl1*^*-/-*^ mice. **(A)** Reduced pericyte coverage of BAT vessels in *Lyl1*^*-/-*^ mice. Left panels: Microscopy images illustrating the severe reduction of pericyte coverage of BAT blood vessels in *Lyl1*^*-/-*^ mice. Scale bar: 30µm. Right panel: Quantification of NG2-positive cell coverage as the ratio between the NG2-positive area and the CD31-positive area. Data are presented as the mean ± SD. **(B)** *Lyl1*^*-/-*^ ingWAT vessels show reduced VE-cadherin recruitment to endothelial cell junctions. FITC-labeled lectin (green) was retro-orbitally injected in 12-week-old WT and *Lyl1*^*-/-*^ mice to visualize blood vessels. After 30 min, ingWAT tissues were collected and 2mm^3^ fragments stained with anti-VE-cadherin antibodies (red). Left panels: Microscopy images illustrating the strong reduction of VE-cadherin staining in *Lyl1*^*-/-*^ compared with WT ingWAT. Scale bar: 30µm. Right panel: quantification of VE-cadherin as the ratio between VE-cadherin staining intensity and lectin area. Data are presented as the mean ± SD. **(C)** Increased leakiness of *Lyl1*^*-/-*^ blood vessels in ingWAT and BAT. Evans blue dye was intravenously injected in 12-week-old WT and *Lyl1*^*-/-*^ mice. After 30min, dye extravasation was measured in ingWAT and BAT, as described in the Methods section. **(D)** *Lyl1* deficiency increases the angiogenic potential of ingWAT stromal vascular particulates (SVPs). Representative images of three experiments. The angiogenic response in each SVP was determined by measuring the length of the growing microtubules with Image J and analyzed with the Mann-Whitney test. Scale bar: 80µm. \*\**P*<0.01; *** P<0.001.

We then isolated stromal vascular particulates (SVPs) [38] from adipose tissues of 12-week-old mice, using a procedure that maintain the native stromal vascular structure while removing all mature adipocytes, and compared their angiogenic potential. After 5 days of culture, we stained SVPs with an anti-CD31 antibody and determined the angiogenic response of each individual SVP culture by measuring the length of the growing microtubules. New endothelial tubes were more numerous and longer in SVP cultures from *Lyl1*^*-/-*^ than WT ingWAT (Figure 6D) and eWAT (Figure S7C) samples. Thus, as observed in tumor and lung vessels [16, 21], the vascular structures of *Lyl1*^*-/-*^ adipose tissues exhibit an immature phenotype, as indicated by the impaired VE-cadherin recruitment at adherens junctions and reduced pericyte coverage. Consequently, adipose blood capillaries are leaky and more prone to angiogenesis in *Lyl1*^*-/-*^ than in WT mice.

### Prerequisite angiogenesis for triggering adipogenesis starts earlier in *Lyl1*^*-/-*^ mice

Sprouting angiogenesis is an essential event to trigger adipogenesis during early postnatal [41] as well as adult [42] adipose tissue development. Given our above observations (Figure 6), we chose to analyze the vascular network in 1-week-old ingWAT and in 17.5 day-postcoitus embryos (E17.5) BAT, i.e, the time points that precede the onset of adipogenic differentiation as above described (Figure 4A). IngWAT depots from six-day-old puppies were double stained with anti-CD31 and anti-NG2 antibodies to visualize vessel structures. Both CD31-positive surface and vessel branching were significantly higher in *Lyl1*^*-/-*^ than WT, revealing that ongoing sprouting angiogenesis was strongly increased in *Lyl1*^*-/-*^ ingWAT (Figure 7A). This was confirmed by the visualization of loosely attached NG2-positive pericytes in *Lyl1*^*-/-*^ ingWAT when compared to the packed and regular coverage of vessels in WT ingWAT (Figure 7B and Figure S7A). Similarly, CD31 staining of E17.5-BAT vessels also showed higher angiogenesis in *Lyl1*^*-/-*^ than WT (Figure 7C). Together, these data demonstrate that in *Lyl1*^*-/-*^ immature adipose tissues, angiogenesis starts earlier causing their premature and faster development.

**Figure 7.**
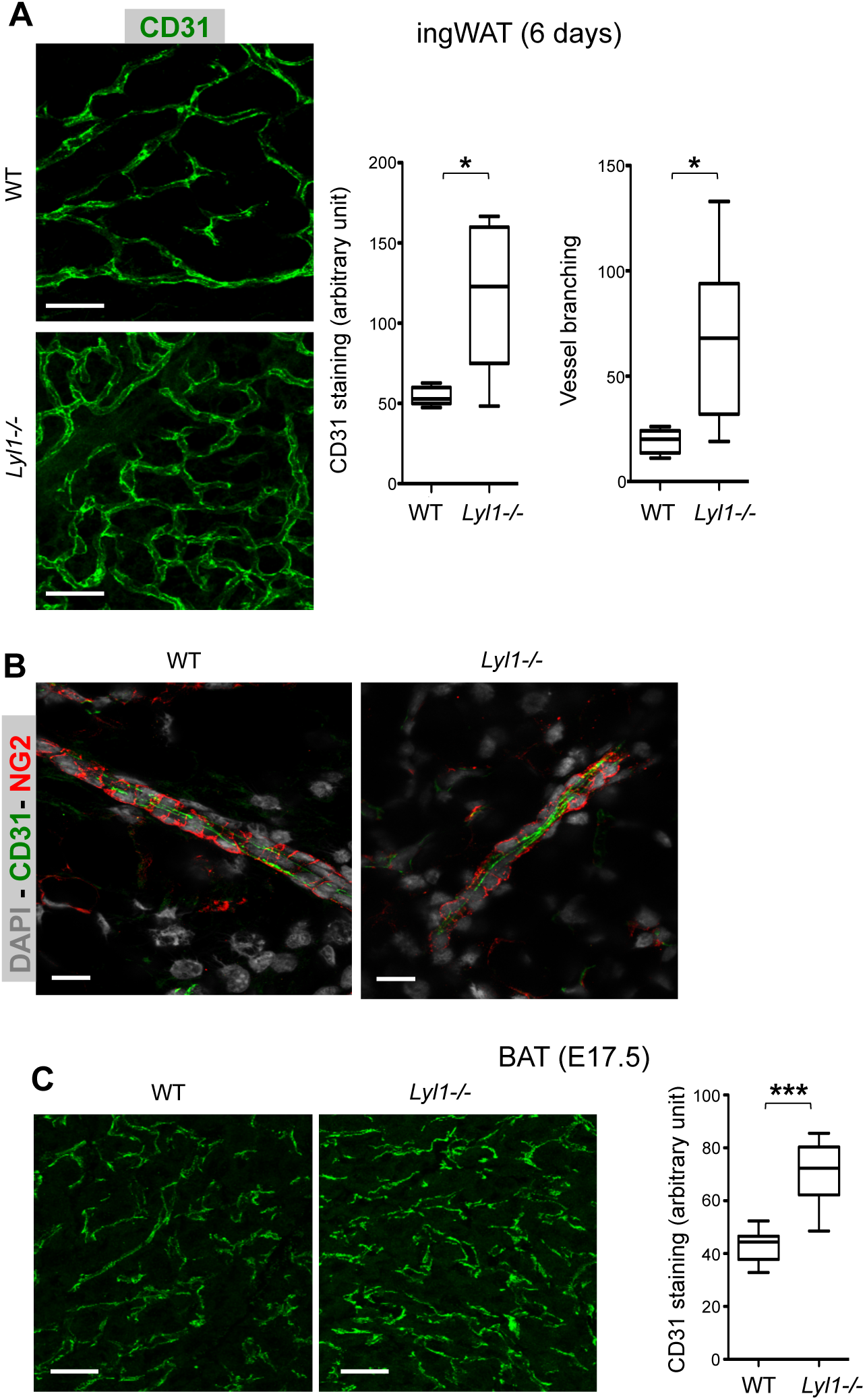
Prerequisite angiogenesis for triggering adipogenesis is earlier in *Lyl1*^*-/-*^ mice. **(A)** Increased angiogenesis in 6day-old *Lyl1*^*-/-*^ ingWAT. Left panels: Microscopy images illustrating the increased vessel development in ingWAT of *Lyl1*^*-/-*^ mice. IngWAT cryosections were stained with antibodies against CD31 (green). Scale bar: 30µm. Right panel: Quantification of CD31 staining and vessel branching. Results are presented as the median (whiskers = min to max) and analyzed with the unpaired *t* test with Welch’s correction. \**P*<0.05. n=5-7mice/genotype. **(B)** NG2+ pericytes (red) are loosely attached to CD31+ EC (green) in *Lyl1*^*-/-*^ ingWAT vessels. Scale bar: 15µm. **(C)** Increased vascular network development in E17.5-old *Lyl1*^*-/-*^ BAT. Left panels: Microscopy images illustrating the increased vessel development in BAT of *Lyl1*^*-/-*^ mice. BAT paraffin-embedded sections were stained with antibodies against CD31 (green). Scale bar: 20µm. Right panel: Quantification of CD31 staining. Results are presented as the median (whiskers = min to max) and analyzed with the unpaired *t* test with Welch’s correction. \*\*\**P*<0.001. n=3-5 embryos/genotype.

## DISCUSSION

The objective of this study was to investigate the mechanisms responsible for the transient obesity observed in young *Lyl1*^-/-^ adult mice. In these animals, overweight is linked to the faster development of adipose tissues, due to the earlier differentiation of immature progenitors. Consequently, in *Lyl1*^*-/-*^ mice, the premature decline of adipogenic potential is not linked to cell-autonomous defects of progenitor cells, but to the reduced availability of stem cells. We found that alterations in the vascular compartment of the adipose niche are responsible for the uncontrolled adipocyte differentiation in *Lyl1*^-/-^ mice.

In *Lyl1*^*-/-*^ mice, mature adipocytes develop earlier in the three fat tissue types. The difference with WT is visible at 1 and 3 weeks of age for BAT and ingWAT, respectively, and at 6 weeks of age for eWAT, in agreement with the development kinetics of the three adipose tissues [43]. Consequently, in young adult *Lyl1*^*-/-*^ mice, adipose tissues exhibit prematurely an aging-like phenotype, such as BAT and ingWAT whitening, as illustrated by lipid droplet enlargement and loss of mitochondrial UCP1 expression [30, 32]. In 5-month-old *Lyl1*^*-/-*^ mice, lipid storage in eWAT starts to decrease, as shown by the smaller adipocytes, concomitantly with decreased expression of several lipogenic genes. In addition, *Lyl1*^*-/-*^ eWAT also expresses higher levels of *Lep* and genes encoding inflammatory molecules, as generally observed in older animals [26, 29-31, 44, 45].

Aging has detrimental consequences on the adipose tissue specific functions (i.e., non-shivering thermogenesis in BAT and ingWAT and lipid storage in eWAT) [32, 46]. In agreement with the lower UCP1 expression and loss of beige fat zones, young adult *Lyl1*^*-/-*^ mice have a lower internal temperature at room temperature. However, this premature aging-like phenotype does not appear to be detrimental because in our animal facility, *Lyl1*^*-/-*^ mice can be maintained for 2 years without major physical problems.

Our data indicate that the early adipose expansion in *Lyl1*^*-/-*^ mice occurs at the adipocyte progenitor levels through non-cell autonomous mechanisms. ASCs and pre-adipocytes isolated from WT and *Lyl1*^*-/-*^ mice show a similar adipogenic potential, in agreement with the absence of *Lyl1* expression in the adipocyte lineage. Conversely, the adipogenic potential of SVFs from *Lyl1*^*-/-*^ BAT and ingWAT declines prematurely due to the reduction in cell number that affects particularly the ASC compartment. As it is the case for several tissues [5, 10, 47-49], the adipose tissue vessel wall represents a reservoir of stem cells that might differentiate into pre-adipocytes and adipocytes. In the adipose vascular niche, ASCs function as perivascular pericytes [38, 50, 51] and it has been suggested that some ASCs may derive from specialized ECs [39]. These studies have highlighted the close relationship between adipocyte formation and endothelium. Indeed, sprouting angiogenesis precedes both post-natal and adult adipose tissue development [13, 41, 42]. As we observed in aortic ring assays [16], the vascular structures from *Lyl1*^*-/-*^ eWAT and ingWAT are more prone to *ex vivo* angiogenesis than WT samples. Correspondingly, a highly developed vascular network is found in *Lyl1*^*-/-*^ immature adipose tissues as early as E17.5 in BAT and 1-week in ingWAT, but not in WT. This excessive angiogenesis, which characterizes *Lyl1*^*-/-*^ vascular structures, is likely to be responsible for the premature and faster development of adipose tissues in *Lyl1*^*-/-*^ mice.

Similarly to *Lyl1*^*-/-*^ lung blood vessels [21], *Lyl1*^*-/-*^ vascular structures in adipose tissues display leakiness due to impaired recruitment of VE-CADHERIN to adherens junctions and poor vessel coverage by pericytes. A recent study demonstrated that according to their permeability properties, blood vessels could support either stem cell quiescence or activation [52]. Indeed, in the hematopoietic stem cell (HSC) niche, less permeable arterial blood vessels maintain HSC quiescence, whereas more permeable sinusoids promote HSC activation and differentiation. Thus, we propose that the immature and unstable vascular structures of *Lyl1*^*-/-*^ adipose tissues prevents ASC maintenance and self-renewal on the vessel wall, thereby accelerating their differentiation into adipocytes and causing premature stem cell depletion. Importantly, the adipose tissue niche might not be the only affected vascular niche in *Lyl1*^*-/-*^ mice. *Lyl1*^*-/-*^ adult bone marrow contains an excess of myeloid cells (our unpublished data) and this could be correlated with the presence of fewer HSCs [17]. Indeed, several studies demonstrated that any HSC niche alteration skews HSC differentiation toward the myeloid lineage [53-56].

In conclusion, we show here that *Lyl1* plays a crucial role in ASC maintenance in a non-cell autonomous manner by controlling the adipose tissue endothelial niche. Our study confirms the major structural role of the vascular niche in coordinating self-renewal or differentiation of stem cells within different tissues. It emphasizes that vascular alterations of the niche may profoundly affect the fate of tissue-residing stem cells, thereby disturbing postnatal tissue homeostasis.

## ACKNOWLEDGEMENTS

We are grateful to the “Réseau des Animaleries de Montpellier” RAM-IBiSA facility and the “Zone d’Expérimentation et de Formation de l’IGMM”-ZEFI facility for animal breeding and experiments. We thank C. Bertrand-Gaday and F. Casas (METAMUS platform) for metabolic phenotyping. We are indebted to the Montpellier Resources Imaging facility for microscopy image acquisition and analysis and to the “Réseau d’Histologie Experimentale de Montpellier”-RHEM platform for expert assistance with histology. AH was supported by a scholarship from The Higher Education Commission (HEC, Pakistan). NP, DM and VP are supported by INSERM.

