## Supplemental figures table and material for "In *Lyl1*^*-/-*^ mice, adipose stem cell vascular niche impairment leads to premature development of fat tissues"

### Figure S1

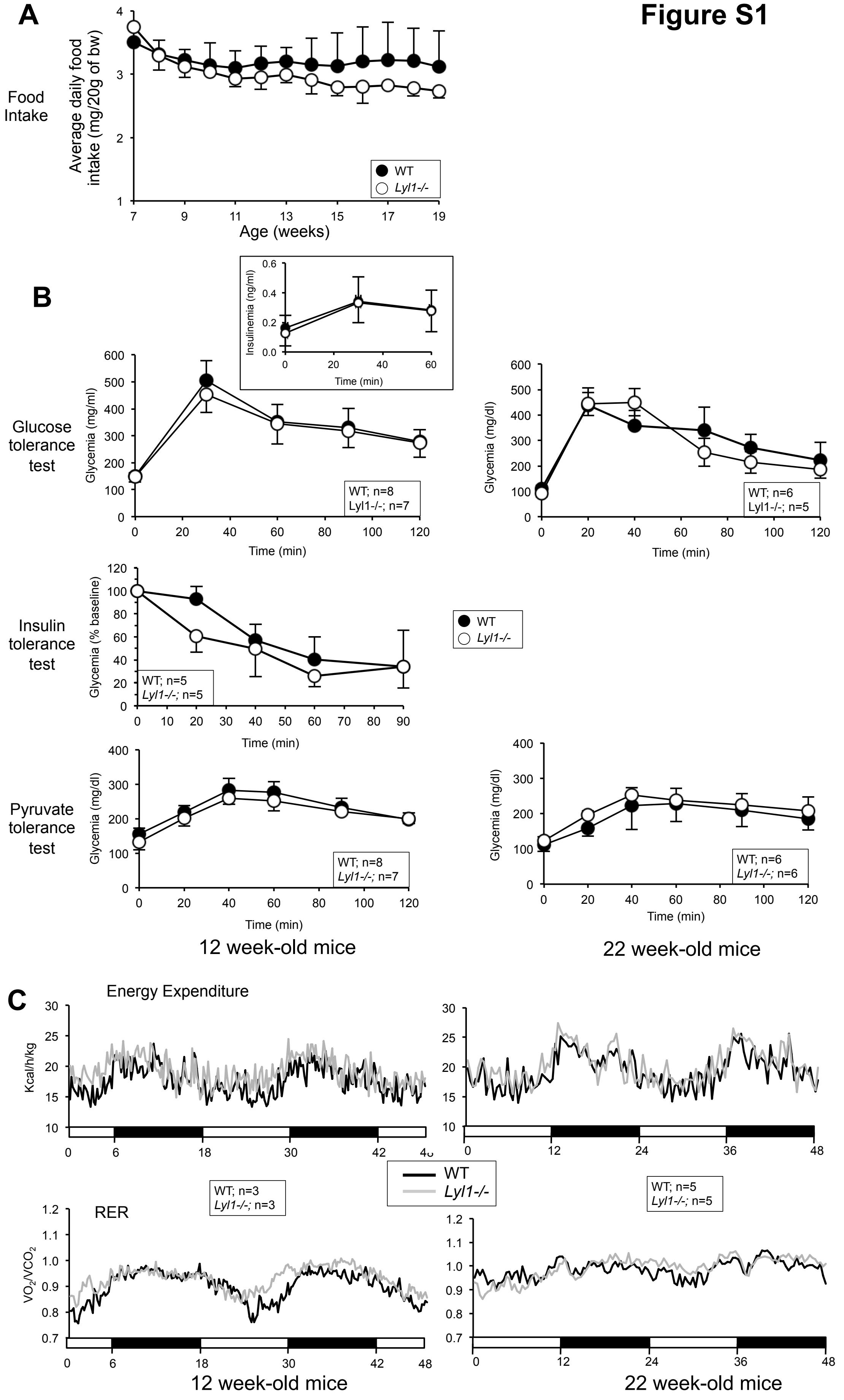
A
Food Intake
Average daily food intake (mg/20g of bw)
Age (weeks)
WT
Lyl1-/-
B
Insulinemia (ng/ml)
Time (min)
Glucose tolerance test
Glycemia (mg/ml)
WT; n=8
Lyl1-/-; n=7
Glycemia (mg/dl)
WT; n=6
Lyl1-/-; n=5
Insulin tolerance test
Glycemia (% baseline)
WT; n=5
Lyl1-/-; n=5
Pyruvate tolerance test
WT; n=8
Lyl1-/-; n=7
WT; n=6
Lyl1-/-; n=6
12 week-old mice
22 week-old mice
C
Energy Expenditure
Kcal/h/kg
WT; n=3
Lyl1-/-; n=3
WT; n=5
Lyl1-/-; n=5
RER
VO2/VCO2
12 week-old mice
22 week-old mice

**Figure S1.** ***Lyl1-/-*** **mice display no obvious metabolic disorders classically linked to obesity.**
**(A)** Food intake is presented as the quantity of ingested food normalized to 20g of body weight. Food was weighted once per week for 12 consecutive weeks for WT (n=5) and *Lyl1*$^{-/-}$ (n=4) mice.
**(B)** For glucose tolerance (GTT) and pyruvate tolerance (PTT) tests, males were fasted overnight and injected intraperitoneally with 2g/kg of glucose or sodium pyruvate, respectively. For insulin resistance (ITT) test, males were fasted 4h and injected intraperitoneally with 0.75 U/kg of insulin. Glucose concentration was measured with a blood glucometer (Accu-Chek, Roche).
**(C)** Male mice were housed in individual cages with a precise control of the temperature and light/dark cycle, with *ad libitum* access to standard chow diet and water for 24h, prior to a 48h period of automated recordings. For each mouse, VO2 and VCO2 were recorded every 13min during the entire experiment. The respiratory exchange ratio (RER) was calculated as the volume of CO2 versus volume of oxygen (VCO2/VO2) ratio.

#### Figure S2

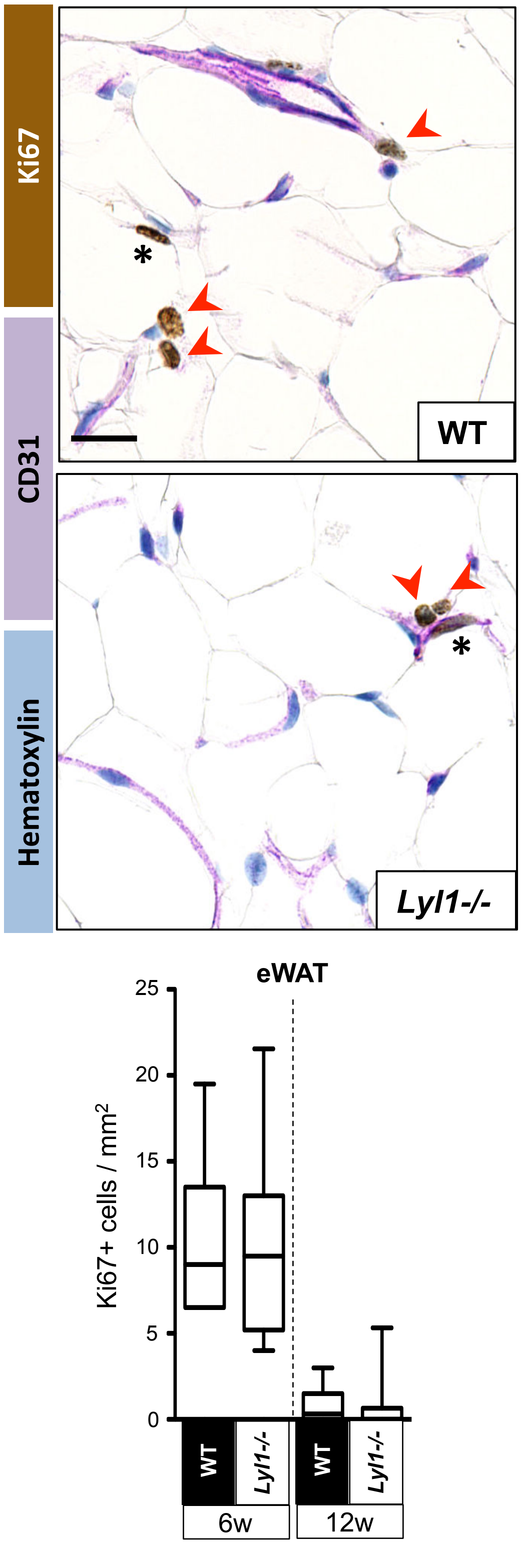


**Figure S2. *Lyl1-/-* and WT adipose tissues display similar proliferative status.**
**Upper panel:** eWAT tissue sections from WT and *Lyl1*$^{-/-}$ mice were co-immunostained for Ki67 proliferative marker (brown) and CD31 (endothelial cells, purple) and images visualized with a NanoZoomer slide scanner controlled by the NDP.view software. Scale bar: 20µm.
**Lower panel:** Ki67+ cells with a round nucleus (red arrow head) were manually counted whereas Ki67+ cells with an elongated nucleus (stars, proliferating endothelial cells) where excluded. For each samples a minimum of 3 areas of 3-5mm$^2$ each were counted. WT (n=4-7 per age) and *Lyl1*$^{-/-}$ (n=4-8 per age).

### Figure S3

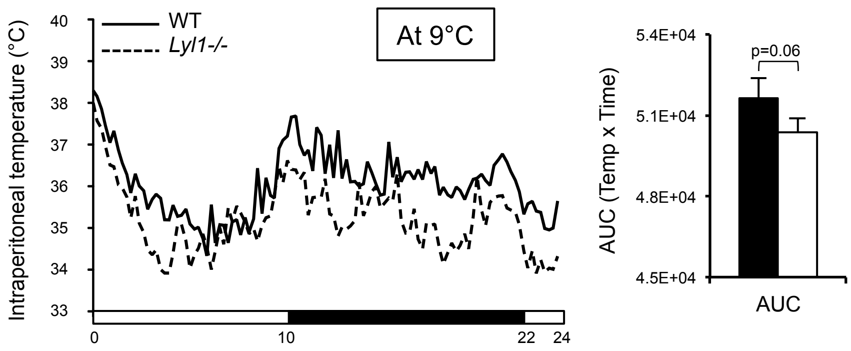


**Figure S3. *Lyl1-/-* mice respond to cold stress as efficiently as WT mice.**
Intraperitoneal temperature was monitored in 12-week-old mice housed individually in metabolic cages with a temperature of 9°C for 24hrs (3 mice for each genotype). The area under the curve (AUC) was calculated using the trapezoidal rule.

### Figure S4

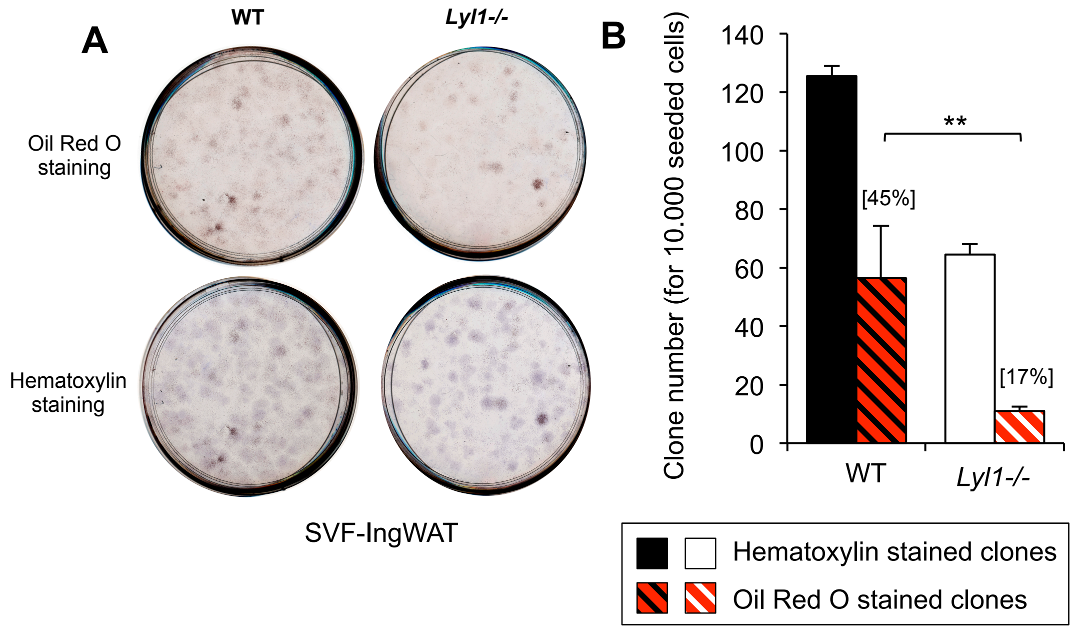


**Figure S4. Lower clonogenic capacity of *Lyl1-/-* SVF-IngWAT compared to WT.**
**(A)** SVF cells from WT and *Lyl1-/-* IngWAT were seeded in equal quantity and when clones reached 50-100 cells adipocyte differentiation was induced with adipogenic cocktail for 2days. After 7days, clones differentiated into adipocytes were stained with Oil RedO while hematoxylin staining revealed the presence of all the clones. Representative image of 3 experiments.
**(B)** Oil RedO stained clones were counted using ImageJ software and their number reported to total number of clones; ***P*<0.01.

### Figure S5

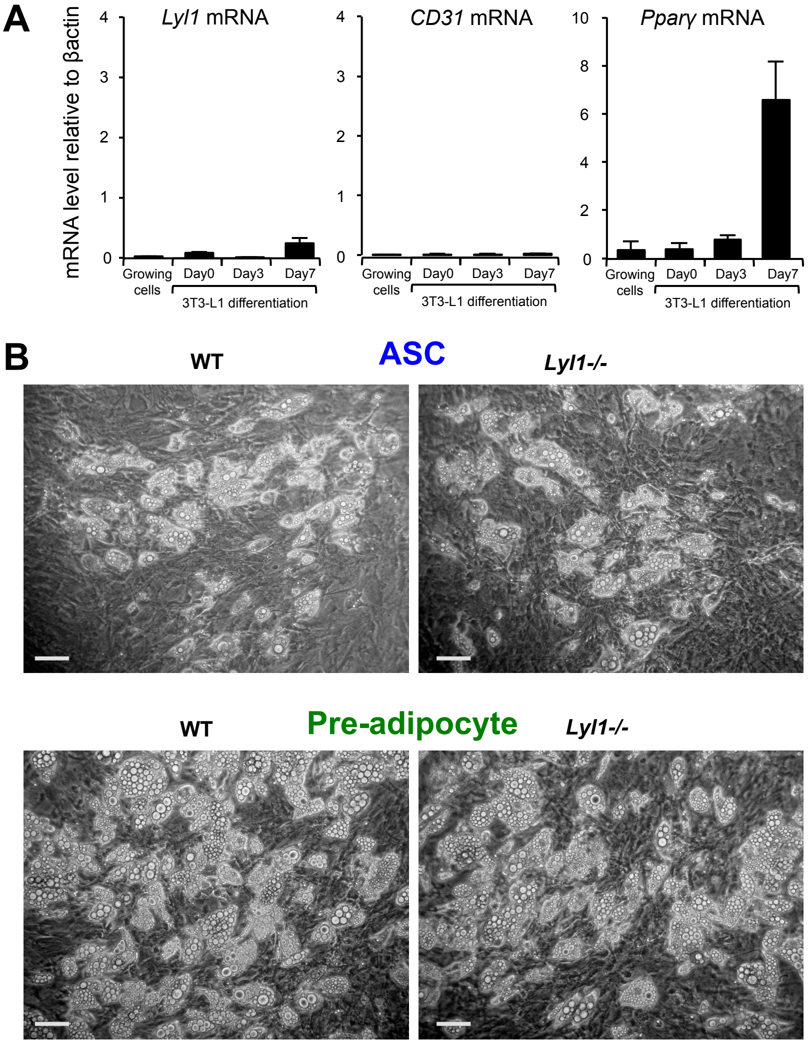


**Figure S5.**

**(A) *Lyl1* is not expressed in 3T3-L1 cells.** Confluent mouse 3T3-L1 cells were incubated with the adipogenic differentiation cocktail and total RNA extracted at day 0, 3 and 7 post-induction. Analysis of *Lyl1*, *CD31* (endothelial cell marker) and *Pparγ* mRNA expression were performed by RT-PCR.

**(B) Isolated ACSs and pre-adipocytes derived from Lyl1-/- mice exhibit similar adipogenic potential as those derived from WT mice**. SVF were prepared from 12-week-old WT and *Lyl1-/-* IngWAT. ASCs (A) and pre-adipocytes (B) were sorted using BD FACS Aria cell sorter. Equal number of cells were plated and induced to differentiate with adipogenic cocktail. Representative images from WT and *Lyl1-/-* 7day-differentiated ASCs (A) and 6-day-differentiated pre-adipocytes (B). Scale bar: 100µm.

Figure S6

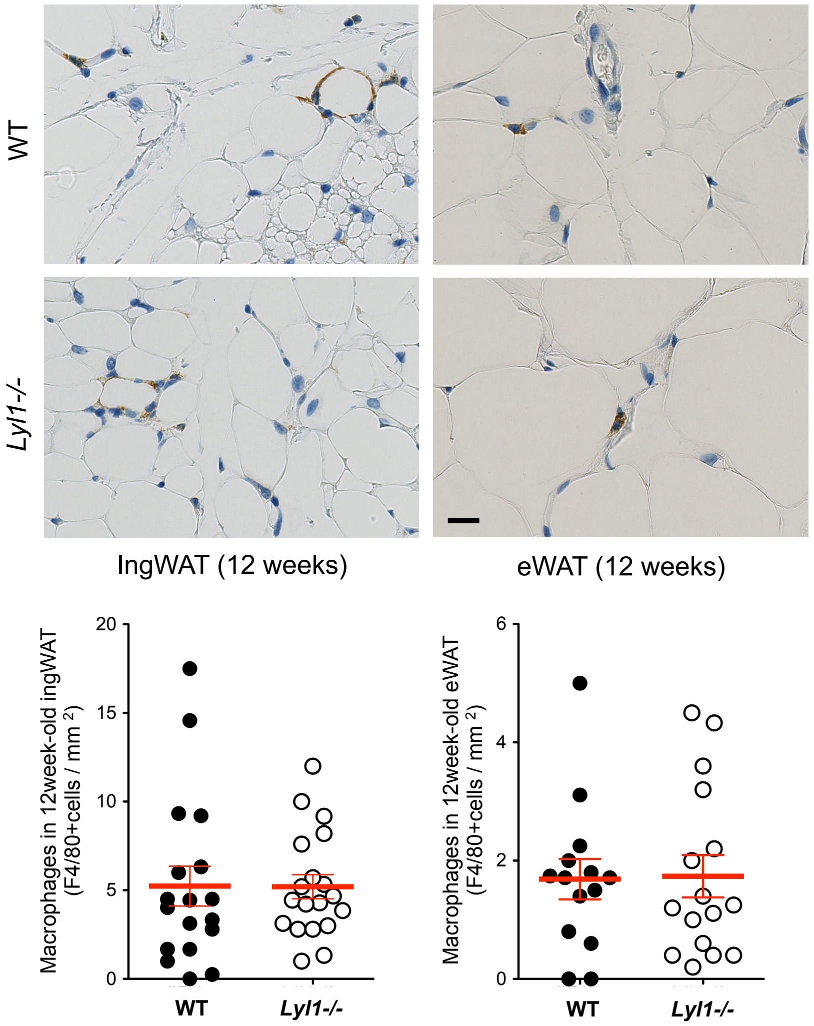


**Figure S6. Equal macrophage infiltration in WT and *Lyl1-/-* adipose tissues.** Macrophages were immuno-stained with anti-F4/80 antibody in paraffin-embedded ingWAT and eWAT sections from 12week-old WT (n=5) and *Lyl1-/-* (n=5) mice. Cells were manually counted in at least 3 areas/mouse tissue visualized with NDP.Viewer software. Scale bar: 20µm.

### Figure S7

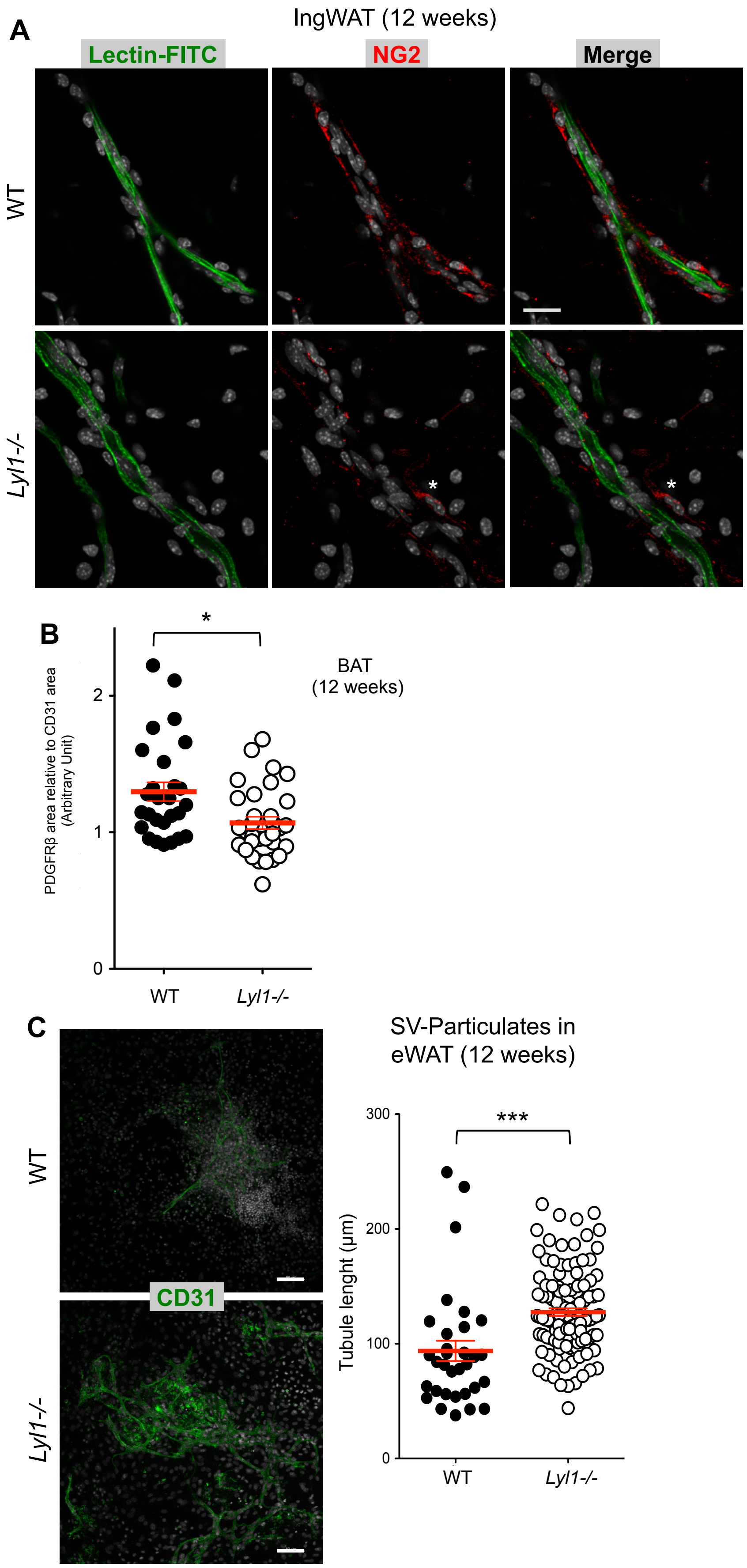
A
IngWAT (12 weeks)
Lectin-FITC
NG2
Merge
WT
Lyl1-/-
*
*
B
*
BAT
(12 weeks)
PDGFRβ area relative to CD31 area
(Arbitrary Unit)
2
1
0
WT
Lyl1-/-
C
SV-Particulates in
eWAT (12 weeks)
WT
CD31
Lyl1-/-
300
200
100
0
***
Tubule lenght (µm)
WT
Lyl1-/-

**Figure S7.**

**(A) *Lyl1-/-* vessels are disorganized.** FITC-labeled Lectin (green) was retro-orbitally injected in 12-week-old WT and *Lyl1-/-* mice to visualize blood vessels. After 30 min, IngWAT tissues were collected and 2mm$^3$ fragments stained with NG2 (red). Perivascular pericytes are less adherent (star) in Lyl1-/- IngWAT.

**(B) Reduced pericyte coverage of BAT vessels in *Lyl1$^{-/-}$* mice.** Quantification of PDGFRβ-positive cell coverage as the ratio between the PDGFRβ-positive area and the CD31-positive area. Data are presented as the mean ± SD. n=8-10 mice/genotype.

**(C) *Lyl1* deficiency increases the angiogenic potential of eWAT stromal vascular particulates (SVPs)**. eWAT samples from 12-week-old WT and *Lyl1$^{-/-}$* mice were mildly digested to maintain the vascular structure integrity. After 5 days of growth, structures were incubated with an antibody against the endothelial cell marker CD31 (green). Representative images of three experiments. Images of microtubule outgrowth from SVPs were captured with a Leica SP5-SMD confocal microscope. The angiogenic response in each SVP was determined by measuring the length of the growing microtubules with Image J and analyzed with the Mann-Whitney test. Scale bar: 80µm. *** P<0.001.

### SUPPLEMENTAL MATERIALS

***Metabolic parameter survey***

For glucose tolerance and pyruvate tolerance tests, 2g/kg of glucose or sodium pyruvate, respectively, were injected intraperitoneally in males after overnight fasting. For insulin resistance testing, 0.75 U/kg of insulin was injected intraperitoneally in males after 4h-fasting. Glucose concentration was measured with a blood glucometer (Accu-Chek, Roche). Oxygen consumption and carbon dioxide production were measured using a Comprehensive Lab Animal Monitoring System (Columbus Instruments, Columbus, OH, USA). Male mice were acclimatized individually in metabolic cages (with precise control of temperature and light/dark cycle) with *ad libitum* access to SCD and water for 24h, prior to a 48h period of automated recordings. For each mouse, $O_2$ volume ($V_{O2}$) and $CO_2$ volume ($V_{CO2}$) were recorded every 13min during the entire experiment. The respiratory exchange ratio was calculated as the $V_{CO2}/V_{O2}$ ratio. The core body temperature was constantly monitored in each mouse in metabolic cages using telemetry equipment and mini-transmitters that were surgically implanted in the abdominal cavity after anesthesia, as recommended by the supplier (Columbus Instruments). Experiments were performed at least 2 weeks after the implant to allow full recovery. For exposure to cold temperatures, non-fasted mice were acclimatized individually in metabolic cages at 22-24°C, prior to their transfer into 9°C pre-cooled cages for 24h.

***RNA preparation and mRNA expression analysis***

Mouse tissues were snap-frozen in liquid nitrogen and kept at -80°C until lysis in TRIzol reagent (Thermo Fischer Scientific, France). Total cell RNA was extracted using the High Pure RNA Isolation Kit (Roche) following the manufacturer's instructions. RNAs were reverse transcribed with a random primer and SuperScript II (Invitrogen). The sequences of the primers used for PCR amplification are listed in Table S1. Gene expression levels were normalized to *Actβ* expression in tissue samples and to *36b4* in isolated adipocyte cells. All results are expressed as mean ± standard deviation (SD).

##### *Immunohistochemical staining*

For immunohistochemistry, 4µm-thick tissue sections were processed as previously described (Pirot et al., 2014), incubated with a sheep anti-UCP1 antibody (kindly provided by D. Ricquier, Paris), for 1h and then biotinylated secondary antibody for 45min. Ki67, CD31 and F4/80 immunohistochemical stainings were performed on the Discovery Ultra Automated IHC staining system using the Ventana DAB Map detection kit. Following deparaffination with Discovery EZ Prep solution at 75°C for 24 minutes, antigen retrieval was performed at 95–100°C for 24 using Discovery CC1 buffer respectively for dual staining Ki67/CD31 or for 15 minutes using Discovery RiboCC buffer for F4/80 staining. Endogenous peroxidase was blocked with Discovery Inhibitor CM for 8 minutes at 37°C. For Ki67/CD31 dual staining, the slides were incubated after rinsing at 37°C for 60 minutes with a rabbit anti-Ki67 antibody (Spring Biosciences, M3064, 1:500), following by a stripping in Discovery CC2 buffer 8min at 100°C then antigen retrieval for 24 minutes at 91°C and an incubation with a rabbit anti-CD31 antibody (Abcam, Ab28364, 1:100) for 60 minutes. Signal enhancement was performed using the Discovery DAB Rabbit OmniMap Kit for Ki67 staining and Discovery Purple Rabbit UltraMap Kit for CD31 staining. For F4/80 staining, the slides were incubated after rinsing at 37°C for 60 minutes with a rat anti-F4/80 antibody (Invitrogen, MF48000, 1:100). Signal enhancement was performed using a rabbit anti-rat IgG (H+L) as the secondary antibody (Thermo Scientific, 31219) and the Discovery DAB Rabbit HQ Kit. Slides were then counterstained for 8 minutes and manually dehydrated before coverslips were added. Colored reactions were visualized with the NanoZoomer slide scanner controlled by the NDP.view software. UCP1 staining was quantified with the Aperio Image Scope software.

##### *Adipogenic differentiation*

SVF cells, ASCs and pre-adipocytes were seeded in 12-well plates and cultured until confluence. After two additional days of culture without bFGF, adipogenesis was triggered with induction medium containing 5µg/ml insulin, 0.5mM 3-isobutyl-1-methylxanthine (IBMX), 1µM dexamethasone and 0.2nM T3 hormone (Sigma-Aldrich, France) for 2 or 3 days. For SVF and ASCs purified from ingWAT, induction medium also included 1µM rosiglitazone.

After induction, cells were grown in adipogenic maintenance medium containing 5µg/ml insulin for the indicated time. For ASCs, 0.5µM dexamethasone was also included.

***Clonogenicity test and Oil Red O staining***

An equal number of cells (20,000 cells/90mm Petri dish) from WT and *Lyl1*$^{-/-}$ ingWAT-SVF was seeded and cultured in DMEM-F12 with Glutamax (Thermo Fischer Scientific, France) supplemented with 10% FBS and 5ng/ml bFGF until 50-cell clones appeared. Adipocyte differentiation was induced as described above for 4 days. After induction, cells were kept in adipogenic maintenance medium containing 5µg/ml insulin for 12 days before fixation in 4% paraformaldehyde. Oil Red O stock solution was prepared by dissolving 0.3 g of Oil Red O (Sigma-Aldrich, France) in 100 mL of isopropanol, stirred overnight, then filtered through 0.2 µm filters and stored at 4°C. Oil Red O working solution was prepared by mixing six parts of Oil Red O stock solution with four parts of water (by volume) and filtered through 0.2 µm filters. For staining, cells were incubated with 60% isopropanol for 5min, with Oil Red O working solution for 10min and then washed gently three times with $H_2O$. Stained clones were imaged using an Epson Perfection V750 PRO scanner and counted with ImageJ.

***Blood vessel immunofluorescence staining***

10µm-thick fixed cryostat sections of BAT from 12-week-old mice were co-incubated with the anti-CD31 (BD Biosciences, 557355), anti-NG2 (Chemicon international, AB5320) and anti-PDGFRβ (Millipore, 06-498-I) antibodies, followed by chicken anti-rat IgG Alexa 488 (Molecular Probes, A21470) and donkey anti-rabbit IgG Alexa 555 (Molecular Probes, A31572). Nuclei were stained with DAPI. For lectin staining, 200µl of Fluorescein Griffonia (Bandeiraea) Simplicifolia Lectin I (0.5mg/ml, Vector Laboratories, FL-1101) were injected in the tail vein of 12-week-old WT and *Lyl1*$^{-/-}$ males, 10min before ingWAT tissue collection. Samples were fixed in 4% paraformaldehyde for 2h, finely cut into 2mmx2mm pieces and incubated with an anti-mouse VE-cadherin antibody (kindly provided by P. Huber, Grenoble, France) at 4°C overnight.

***Extravasation of albumin-Evans blue within tissues***

20mg/kg of Evans blue dye (EBD, SIGMA) was injected in the tail vein. Thirty min after injection, animals were anesthetized by i.p. injection of 2mg/kg xylazine (Rompun 2%; Bayer Pharma, France) and 50mg/kg ketamine (Imalgène 500; Merial, France) and the chest was opened. Mice were transcardially perfused with 50ml of PBS/5mM EDTA to remove excess EBD. BAT, eWAT and ingWAT were excised, dried, weighed and placed in formamide at 60°C for 36h. EBD in supernatants was quantified by spectrophotometry at 630 nm.

***Whole-mount confocal microscopy***

Approximately 2mmx2mm ingWAT depots were dissected from 1-week-old male mice and fixed at -20°C in acetone-methanol (30-70%) for 20min. Non-specific antibody binding was blocked by incubation in PBS with 5% normal horse serum and 3% TritonX100 for 5h. Samples were incubated with primary anti-CD31 and -NG2 antibodies at 4°C, under agitation for 40h, and then with chicken anti-rat IgG Alexa 488 and donkey anti-rabbit IgG Alexa 555 for 5h. Nuclei were stained with DAPI. Images were taken on a Leica SP5-SMD confocal microscope and analyzed with the Imaris software.

**Table S1**: Sequences of the *Mus Musculus* primers used in RT-qPCR.

| Gene | Forward Primer | Reverse Primer |
|---|---|---|
| *36b4* | TCGCTCCGAGGGAAGGCCGTGGT | GCCCGAGCAGCAGCTGGCACCTTA |
| *AdRβ3* | AGCAGACAGGGACAGAGGGGTTGCCT | TGGAGGGTGGAGAGGGGCGTCCT |
| *Actinβ* | TCCTGGCCTCACTGTCCAC | GTCCGCCTAGAAGCACTTGC |
| *C/ebpα* | TGCGCGGGCGCGAGCCAGTT | GGGCCGCGGCTCCACCTCGT |
| *Cidea* | GGGAGCCCTCATCAGGCCCCTGACAT | TGCTGGCCATCACCCCACGCCG |
| *Dio2* | AGCCGCTCCAAGTCCACTCGCGG | CCAGTGGGCGCTCTGCACTGGCA |
| *Fabp4* | AGCACCCTCCTGTGCTGCAGCCTT | TCCTTGTGGCAAAGCCCACTCCCACTT |
| *Il6* | CTGGTCTTCTGGAGTACCATAGCTACCTGG | AGCCACTCCTTCTGTGACTCCAGCTT |
| *Klf4* | GCACACCTGCGAACTCACACAGGCGA | TCTGGCACTGAAAGGGCCGGTGCC |
| *Lepl* | ACACACGCAGTCGGTATCCGCCAAGCA | AGGCTCTCTGGCTTCTGCAGGCCACT |
| *Lyl1* | TGCTCAACCCGCTCCTGACT | AGCCACTGCAAGTAGCCTGT |
| *Pgc-1a* | ACCGCAGTCGCAACATGCTCAAGCCAA | ACCGGGCCCTCTTGGTTGGCGG |
| *Plin* | GTTACAGCCCTGCCCAACCC | GCCTGGGAAGCGGCACATAG |
| *Pparγ* | AGTGGAGACCGCCCAGGCTTGCTGA | TCCTGGAGCAGGGGGTGAAGGCTCA |
| *Srebp1* | TGCCCACCCCTGCCCTGCACA | CCAGGGTCTGCAGGGGACCTGCCT |
| *Tnfα* | GCCTGTAGCCCACGTCGTAGCAAACCA | AGGGCGTTGGCGCGCTGGCT |
| *Ucp1* | GCCAAAGTCCGCCTTCAGAT | TGATTTGCCTCTGAATGCCC |
| *Zfp423* | TGGGCGGTACCTTCAAGTGCCCCGT | GCGTGCTGGCTCATCGTGTGGTTCTGC |
